# Selective Sweep at a QTL in a Randomly Fluctuating Environment

**DOI:** 10.1101/752873

**Authors:** Luis-Miguel Chevin

**Author notes:** UMR 5175, 1919 Route de Mende, 34293 Montpellier Cedex 5, France.

## Abstract

Adaptation is mediated by phenotypic traits that are often near continuous, and undergo selective pressures that may change with the environment. The dynamics of allelic frequencies at underlying quantitative trait loci (QTL) depend on their own phenotypic effects, but also possibly on other polymorphic loci affecting the same trait, and on environmental change driving phenotypic selection. Most environments include a substantial component of random noise, characterized by both its magnitude and its temporal autocorrelation, which sets the timescale of environmental predictability. I investigate the dynamics of a mutation affecting a quantitative trait in an autocorrelated stochastic environment that causes random fluctuations of an optimum phenotype. The trait under selection may also exhibit background polygenic variance caused by many polymorphic loci of small effects elsewhere in the genome. In addition, the mutation at the QTL may affect phenotypic plasticity, the phenotypic response of given genotype to its environment of development or expression. Stochastic environmental fluctuations increases the variance of the evolutionary process, with consequences for the probability of a complete sweep at the QTL. Background polygenic variation critically alters this process, by setting an upper limit to stochastic variance of population genetics at the QTL. For a plasticity QTL, stochastic fluctuations also influences the expected selection coefficient, and alleles with the same expected trajectory can have very different stochastic variances. Finally, a mutation may be favored through its effect on plasticity despite causing a systematic mismatch with optimum, which is compensated by evolution of the mean background phenotype.

## Introduction

The advent of population genomics and next-generation sequencing has fostered the hope that the search for molecular signatures of adaptation would reach a new era, wherein the recent evolutionary history of a species would be inferred precisely and somewhat exhaustively, and fine details of the genetics of adaptation would be revealed (Stapley *et al.* 2010). Despite undisputable successes, the picture that has emerged in the last decade is more complex. First, the importance of polygenic variation in adaptation has been re-evaluated based on theoretical and empirical arguments (Chevin and Hospital 2008; Pavlidis *et al.* 2008; Pritchard *et al.* 2010; Rockman 2012; Jain and Stephan 2017; Stetter *et al.* 2018; Höllinger *et al.* 2019), and methods have been designed to detect subtle frequency changes at multiple loci that may jointly cause substantial phenotypic evolution (Turchin *et al.* 2012; Berg and Coop 2014; Stephan 2016; Wellenreuther and Hansson 2016; Racimo *et al.* 2018; Josephs *et al.* 2019). Consistent with (but not limited to) polygenic adaptation is the idea that mutations contributing to adaptive evolution do not necessarily start sweeping when they arise in the population, but may instead segregate for some time in the population and contribute to standing genetic variation, before they become selected as the environment changes (Barrett and Schluter 2008; Kopp and Hermisson 2009; Matuszewski *et al.* 2015; Jain and Stephan 2017). After the factors governing such “soft sweeps” and their influence on neutral polymorphism have been characterized (Hermisson and Pennings 2005; Przeworski *et al.* 2005), the debate has shifted to their putative prevalence in molecular data, and perhaps more importantly to their contribution to adaptive evolution (Jensen 2014; Garud *et al.* 2015; Hermisson and Pennings 2017).

Another line of complexity in the search for molecular footprints of adaptation comes from temporal variation in selection. The classical hitchhiking model (Maynard-Smith and Haigh 1974; Stephan *et al.* 1992) posits a constant selection coefficient without specifying its origin. Some models have gone a step further by explicitly including a phenotype under selection, and have shown that even in a constant environment, selection at a given locus may change over the course of a selective sweep, as the mean phenotype in the background evolves through the effects of other polymorphic loci, in a form of whole-genome epistasis mediated by the phenotype (Lande 1983; Chevin and Hospital 2008; Matuszewski *et al.* 2015). In addition, selection is likely to vary in time because of a changing environment. Most environments exhibit substantial fluctuations over time, beyond any trend or large shifts (Stocker *et al.* 2013). These fluctuations are likely to affect natural selection, which emerges from an interaction of the phenotype of an organism with its environment. Interestingly, one of the first attempts to measure selection through time in the wild revealed substantial fluctuations in strength and magnitude (Fisher and Ford 1947), spurring a heated debate about the relative importance of drift versus selection in evolution, and setting the stage for the neutralist-selectionist debate (Wright 1948; Kimura 1968; Yamazaki and Maruyama 1972; Gillespie 1977). Other iconic examples of adaptive evolution also show clear evidence for fluctuating selection (Lynch 1987; Grant and Grant 2002; Bell 2010; Bergland *et al.* 2014; Nosil *et al.* 2018), suggesting that selection *in natura* is rarely purely directional, but instead often includes some component of temporal fluctuations. Part of these fluctuations involve deterministic, periodic cycles, such as seasonal genomic changes in fruit flies (Bergland *et al.* 2014), but random environmental variation also certainly plays a substantial role. In fact, virtually all natural environments exhibit some stochastic noise, characterized not only by its magnitude but also by its temporal autocorrelation, which determines the average speed of fluctuations and the time scale of environmental predictability (Halley 1996; Vasseur and Yodzis 2004). The influence of such environmental noise on natural populations is attested notably by stochasticity in population dynamics (Lande *et al.* 2003; Ovaskainen and Meerson 2010), and natural selection at the phenotypic level has also been estimated as a stochastic process in a few case studies (Engen *et al.* 2012; Chevin *et al.* 2015; Gamelon *et al.* 2018).

Population genetics theory has a long history of investigating randomly fluctuating selection. In particular, Wright (1948) used diffusion theory to derive the stationary distribution of allelic frequencies in a stochastic environment, which was later extended to find the probability of quasi-fixation in an infinite population (Kimura 1954), and of fixation in a finite population (Ohta 1972). This topic gained prominence during the neutralist-selection debate, where the relative influences of genetic drift vs a fluctuating environment as alternative sources of stochasticity in population genetics was strongly debated with respect to the maintenance of polymorphism and molecular heterozygosity (Nei 1971; Gillespie 1973, 1977, 1979, 1991; Nei and Yokoyama 1976; Takahata and Kimura 1979), a question that remains disputed in the genomics era (Mustonen and Lassig 2007, 2010; Miura *et al.* 2013). Another line of research has asked what is the expected relative fitness of a genotype/phenotype in a fluctuating environment, and whether Wright’s (1937) adaptive landscape could be extended to this context (Lande 2007; Lande *et al.* 2009).

However, this literature is mostly disconnected from the literature on adaptation of quantitative traits to a randomly changing environment (Bull 1987; Lande and Shannon 1996; Chevin 2013; Tufto 2015). Even in work that investigates fluctuating selection both at a single locus and on a quantitative trait (e.g. Lande 2007), the selection coefficient at the single locus is often postulated ad hoc, rather than stemming from its effect on a trait under selection. Connallon and Clark (2015) recently investigated the influence of a randomly fluctuating optimum phenotype on the distribution of fitness effects of mutations affecting a trait, but they assumed non-autocorrelated fluctuations, and did not derive the stochastic variance of the population genetic process, which is important driver of probabilities of (quasi-)fixation (Kimura 1954; Ohta 1972). They also did not consider fitness epistasis caused by evolution of the mean background phenotype. Lastly, this work has largely overlooked possible mutation effects on phenotypic plasticity, the phenotypic response of a given genotype to its environment of development or expression (Schlichting and Pigliucci 1998; West-Eberhard 2003), which is expected to evolve in environments that fluctuate with some predictability (Gavrilets and Scheiner 1993a; Lande 2009; Tufto 2015). Instead, Connallon and Clark (2015) included a form of environmental noise in phenotypic expression that is similar to bet hedging (Svardal *et al.* 2011; Tufto 2015).

I here extend a model that combines population and quantitative genetics (Lande 1983; Chevin and Hospital 2008) to the context of an autocorrelated random environment causing movements of an optimum phenotype, to ask: What is the distribution of allelic frequencies at a QTL in a stochastic environment? How does it depend on whether a mutation is segregating alone, or instead affects a quantitative trait with polygenic background variation? How does environmental stochasticity affect the probability of a complete sweep at the QTL, and the resulting genetic architecture of the trait? And how are these effects altered when the mutation affects phenotypic plasticity?

## Model

### Fluctuating selection

The core assumption of the model is that adaptation is mediated by a continuous, quantitative trait undergoing stabilizing selection towards an optimum phenotype that moves in response to the environment, as typical in models of adaptation to a changing environment (reviewed by Kopp and Matuszewski 2014). More precisely, the expected number of offspring in the next generation (assuming discrete non-overlapping generations) of individuals with phenotype *z* is

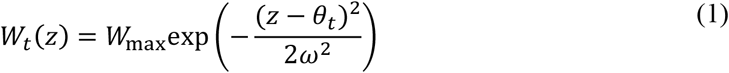

where *θ*_*t*_ is the optimum phenotype at generation *t*, and *ω* is the width of the fitness peak, which determines the strength of stabilizing selection. The height of the fitness peak *W*_max_ may affect demography but not evolution, as it is independent of the phenotype.

In line with other models of adaptation to changing environments (Kopp and Matuszewski 2014), I assume that the environment causes movement of the optimum phenotype, but does not affect the width of the fitness function. The environment undergoes stationary random fluctuations, which may be combined initially to a major, deterministic environmental shift of the mean environment. The stochastic component of variation in the optimum is assumed to be autocorrelated, in the form of a first-order autoregressive process (AR1) with stationary variance 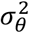 and autocorrelation *ρ* over unit time step (one generation). This is one of the simplest forms of autocorrelated continuous process: it is Markovian (memory over one time step only), leading to an exponentially decaying autocorrelation function with half-time *T*_half_ = −ln(2)/ln(*ρ*) generations.

### Genetics

For simplicity, I base the argument on a haploid model, but much of the findings extend to diploids, with a few additional complications such as over-dominance caused by selection towards an optimum (Barton 2001; Sellis *et al.* 2011). I focus on a mutation at a locus affecting the quantitative trait – i.e., a quantitative trait locus, or QTL -, with additive haploid effect *α* on the trait. More precisely, I consider a bi-allelic QTL, with mean phenotype *m* for the wild-type (ancestral) allele, in frequency *q* = 1 − *p*, and *m* + *α* for the mutant (derived) allele, in frequency *p*. We are not interested here in the origin and initial spread of the mutation from initially very low, drift-dominated frequencies. Investigating this would require extending theory of fixation probabilities in changing environments (Uecker and Hermisson 2011) to include environmental stochasticity, which is beyond the scope of this work. Instead, the focus is here on adaptation from standing genetic variation, and the aim will be to track the evolutionary trajectory of a focal mutation at a bi-allelic locus, starting from a low initial frequency *p*_0_ where most of frequency change can be attributed to selection. We will briefly address the influence of drift at the end of the analysis.

Two types of genetic scenarios will be contrasted. In the “monomorphic background” scenario, no other polymorphic locus affects the quantitative trait when the focal mutation is segregating at the QTL. This corresponds to a form of strong selection weak mutation approximation (SSWM Gillespie 1983, 1991). This scenario requires no further assumption about the reproduction system (sexual or asexual). In the opposite “polygenic background” scenario, variation in the trait is assumed to be caused by a large number of weak-effect loci (or “minor genes”), in addition to the effect of the QTL (or “major gene”). Sexual reproduction is assumed, with fertilization closely followed by meiosis over a short diploid phase where selection can be neglected. I further assume that minor genes are unlinked among themselves and with the major gene, such that the genotypic background has a similar distribution for all alleles at the major gene. Following standard quantitative genetics (Falconer and MacKay 1996; Lynch and Walsh 1998), I assume that additive genetic values in the background are normally distributed, with mean phenotype *m* and additive genetic variance *G*, and that phenotypes also include a residual component of variation independent from genotype, with mean 0 and variance *V*_*e*_. This model of major gene and polygenes, which takes its roots in Fisher’s (1918) foundational paper for quantitative genetics, has been analyzed for evolutionary genetics by Lande (1983), and later used to investigate selective sweeps at a QTL in constant environment or following an abrupt environmental shift by Chevin and Hospital (2008). I here extend this work to a randomly changing environment.

### Phenotypic plasticity

I also investigate the case where both the mean background phenotype and the QTL effect may respond to the environment, *via* phenotypic plasticity. Let *ɛ* be a normally distributed environmental variable (e.g. temperature, humidity…) with mean 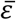 and variance 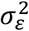, which affects the development or expression of the trait. Assuming a linear reaction norm for simplicity, the mean background phenotype is

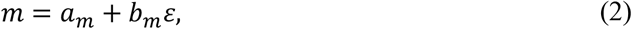

where *b*_*m*_ is the slope of reaction norm, which quantifies phenotypic plasticity, and the intercept *a*_*m*_ is the trait value in a reference environment where *ɛ* = 0 by convention. I neglect evolution of plasticity in the background for simplicity, and therefore assume that *b*_*m*_ is a constant, while *a*_*m*_ is a polygenic trait with additive genetic variance *G* as before. The additive effect of the mutation at the QTL is also phenotypically plastic, such that

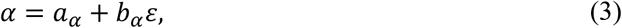

with *b*_*α*_ the additive increase in plasticity caused by the mutation at the QTL, and *a*_*α*_ the additive effect on the trait in the reference environment.

The environment of development partly predicts changes of the optimum phenotype for selection, such that

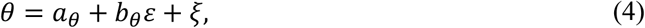

where *ξ* is normal deviate independent from *ɛ*, with mean 0 and variance 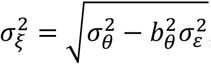, such that the variance of optimum remains 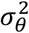. Note that eq. (4) does not necessarily imply a causal relationship between *ɛ* and *θ*, because selection occurs after development/expression of the plastic phenotype and is thus likely to be influenced by a later environment (Gavrilets and Scheiner 1993a; Lande 2009). In fact, the optimum may even respond to other environmental variables than *ɛ*, which jointly constitute the cause of selection (Wade and Kalisz 1990; MacColl 2011), but can be partly predicted by *ɛ* upon development. In this case *b*_*θ*_ is the product of the regression slope of the optimum on the causal environment for selection, times the regression slope of this causal environment on the environment of development *ɛ* (de Jong 1990; Gavrilets and Scheiner 1993a; Chevin and Lande 2015). When the same environmental variable affects development and selection but at different times, then the latter regression slope is simply the autocorrelation of the environment between development and selection within a generation (Lande 2009; Michel *et al.* 2014).

### Evolutionary dynamics

Lande (1983) has shown that the joint dynamics of a major gene and normally distributed polygenes in response to selection are governed by a couple of equations that are remarkably identical to their counterpart without polygenes and without a major gene, respectively. In other words, Wright’s (1937) fitness landscape for genes and Lande’s (1976) fitness landscape for quantitative traits jointly apply in the context of major gene combined with polygene. For a haploid sexual population, the recursions for the allelic frequency *p* of the mutation at the major gene and for the mean phenotype *m* in the polygenic background are then

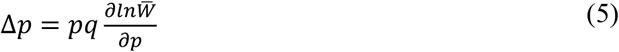

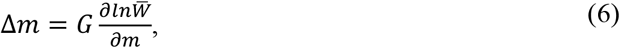

where the partial derivatives are selection gradients on allelic frequency and mean phenotype, respectively (Wright 1937; Lande 1976).

With selection towards an optimum as modeled in equation (1), and an overall phenotype distribution that is a mixtures of two Gaussians with same variance *G* + *V*_*e*_ and modes separated by the effect of the major gene *α*, the mean fitness in the population is

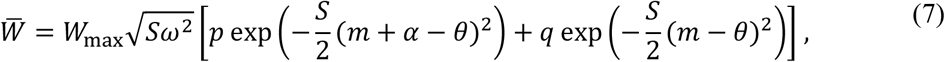

where 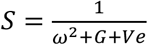 is the strength of stabilizing selection. Combining eqs (6) and (7), the selection gradient on the mean background phenotype is

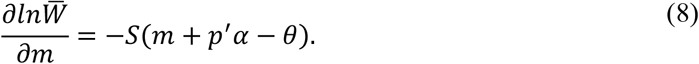

As in classical models of moving optimum for quantitative traits (Lande 1976; Kopp and Hermisson 2007), directional selection on the trait is proportional to the deviation of the mean phenotype from the optimum, multiplied by the strength of stabilizing selection, which is larger when the fitness peak is narrower. However here, the overall mean phenotype depends on *p*′, the frequency after selection of the mutation at the QTL. This causes a coupling of dynamics in the background and at the QTL.

For the dynamics at the QTL it will be convenient to focus on the logit allelic frequency of the mutation, *ψ* = ln(*p*/*q*). With a constant selection coefficient *s* as assumed in classical models of selective sweeps, *ψ* would increase linearly in time with slope *s* (Stephan *et al.* 1992), while *ψ* changes non linearly in time even in a constant environment if the mutation is dominant/recessive (Teshima and Przeworski 2006), or if it affects a quantitative trait with polygenic background variation as assumed here (Chevin and Hospital 2008). Combining eqs. (5) and (7), after some simple algebra the recursion for *ψ* over one generation of selection is

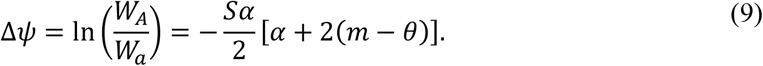

Note that Δ*ψ* is a measure of the selection coefficient *s* for this generation (Chevin 2011). In a constant environment where *θ*_*t*_ = *θ* for all *t*, the system admits two stable equilibria with fixation at the QTL,

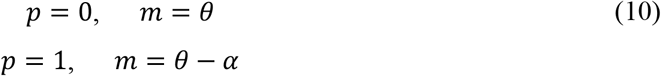

and one unstable internal equilibrium

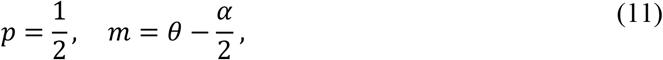

in line with previous analysis of the diploid version of this model (Lande 1983). Note that the mean background phenotype evolves to compensate for the effect of the major gene, such that the overall mean phenotype is at the optimum in all three equilibria, *m* + *pα* = *θ*.

### Approximation for weak fluctuating selection at QTL

The full model with coupled dynamics at the major gene and background polygenes can be used for numerical recursions, but to make further analytical progress, I rely on an approximation of this model that neglects the influence of the QTL on the background mean phenotype, as in previous analysis in a constant environment (Chevin and Hospital 2008). In a randomly fluctuating environment, this approximation consists of assuming that selection at the QTL is sufficiently weak that its contribution to fluctuating selection on the mean background phenotype can be neglected, such that variance in the directional selection gradient is proportional to

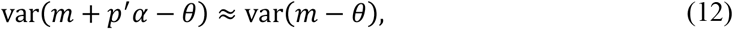

and similarly for its covariance across generations.

### Simulations

The mathematical analysis of this model is complemented by population-based simulations under a randomly fluctuating optimum. These simulations are based on recursions of equations (5–7), assuming a constant additive genetic variance *G* in the background. In each simulation, the optimum is initially drawn from an normal distribution with mean 0 and variance 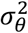, and optima in subsequent generations are drawn using 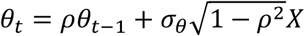, where *X* is a standard normal deviate, such that *θ* has stationary variance 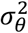 and autocorrelation *ρ* as required. In simulations with phenotypic plasticity, the environment of development is drawn retrospectively from the optimum, using 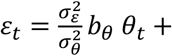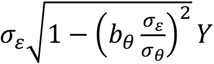, where *Y* is drawn from a standard normal, such that *ɛ* has variance 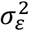 and the regression slope of *θ* on *ɛ* is *b*_*θ*_, as required (eq. 4). In simulations with background genetic variance, the system is left to evolve for 500 generations, to allow the mean background phenotype to reach a stationary distribution with respect to the fluctuating environment. The initial frequency at the QTL is set then to *p*_0_, and the mean optimum is shift by *m*_0_ relative to the expected background mean phenotype. To simulate random genetic drift, the allelic frequency at the QTL in the next generation is drawn randomly from a binomial distribution with parameters *N*_*e*_ (the effective population size) and *p*′ (the expected frequency after selection in the current generation), consistent with a haploid Wright-Fisher population (Crow and Kimura 1970). Similarly for the mean background, genetic drift was simulated by drawing the mean phenotype in the next generation from a normal distribution with mean the expected mean background phenotype after selection, and variance *G*/*N*_*e*_ (Lande 1976).

### Data availability

A Mathematica notebook including code for simulations is available from a FigShare repository.

## Results

We are interested in fluctuating selection at a gene affecting a quantitative trait (or QTL) exposed to a randomly moving optimum phenotype. The stochastic population genetics at the QTL will be analyzed on the logit scale *ψ* = ln(*p*/*q*) for mathematical convenience (as in, e.g., Kimura 1954; Gillespie 1991), and also because this directly relates to empirical measurements (Chevin 2011; Gallet et al. 2012; see also Discussion). From equation (9), *t* generations after starting from an initial logit frequency *ψ*_0_, we have

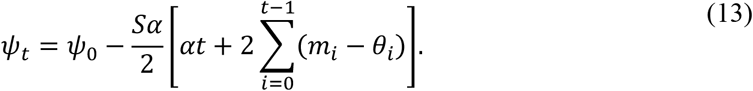

The first term in brackets increases linearly with time, and corresponds to a component of selection that only depends on the phenotypic effect of the mutation and the strength of selection on the trait, but not on the background phenotype or the environment. All the influence of the fluctuating environment and background phenotype arises through the sum (second term in brackets), which shows that the influences of all past maladaptations (deviations of the mean phenotype from the optimum) weigh equally in their contribution to population genetics over time. In a stochastic environment, this means that a chance event causing a large deviation from the optimum can have persistent effects on genetic change. This occurs here because selection is assumed to be frequency independent; with frequency-dependent selection, non-linear dynamics could instead rapidly erase memory of past environments and maladaptation, as occurs for population dynamics with density dependence (Chevin *et al.* 2017).

The optimum phenotype is assumed to follow a Gaussian process. In most contexts we will investigate, this causes the population genetics at the QTL to also follow a Gaussian process on the logit scale, such that *ψ* has a Gaussian distribution at any time. A Gaussian distribution of logit allelic frequency was also found in phenomenological models without an explicit phenotype, where selection coefficients were assumed to undergo a Gaussian process (Kimura 1954; Gillespie 1991, p.149). The reason for this correspondence is that *ψ* is linear in phenotypic mismatches with optimum in eq. (13), and these mismatches themselves follow a Gaussian process (i) in the absence of background polygenic variation; and (ii) with background polygenic variation, as long as evolution of the mean background is little affected by the QTL, such that *m* + *p*′*α* − *θ* ≈ *m* − *θ*. When these assumptions hold, the distribution of allelic frequencies in a stochastic environment can be summarized by their mean and variance on the logit scale, E_*ψ*_ and 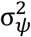. A simple transformation can then be used to retrieve the distribution of allelic frequencies, following Gillespie (1991, p.149),

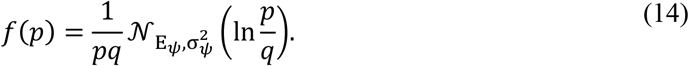

where 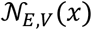 is the density of a normal distribution with mean *E* and variance *V* evaluated at *x*. This transformation is illustrated in **Figure 1**.

**Figure 1:**
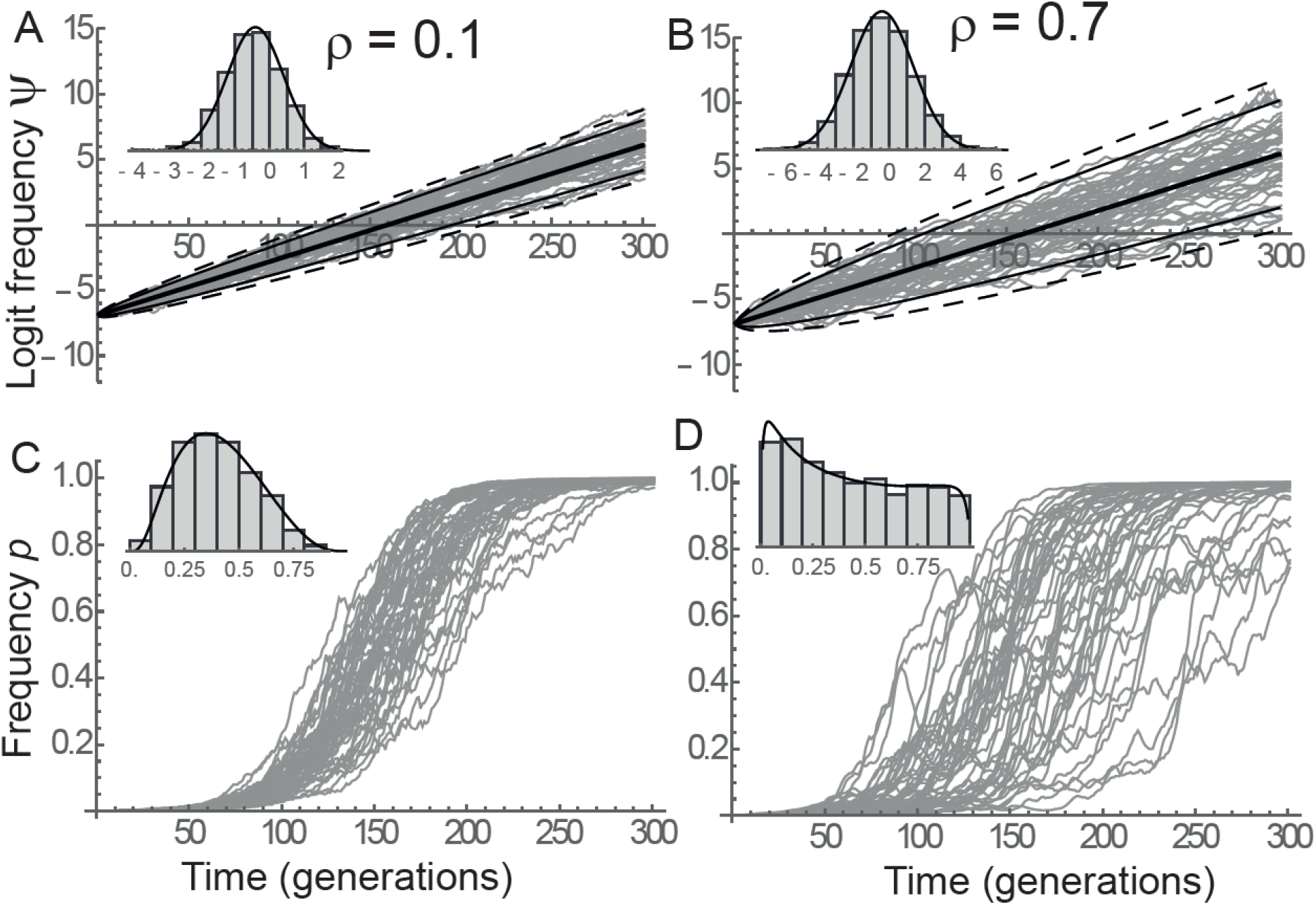
Fluctuating selection at a QTL in a monomorphic genetic background. The dynamics of logit allelic frequency *ψ* (A-B) and allelic frequency *p* (C-D) are shown as gray lines for 50 simulations with low (*ρ* = 0.1, left) or high (*ρ* = 0.7, right) positive autocorrelation in the optimum. Panels A-B also show percentiles from the predicted normal distribution with mean and variance provided by eqs. (15) and (17), respectively: 50% (median) in thick, 5% and 95% in thin, and 1% and 99% in dashed lines. Insets show distributions at generation 150, where histograms are computed from 1000 simulations, while the solid black line is the predicted density based on the moments in eqs. (15) and (17) for A-B, and their transformation using eq. (14) for C-D. Parameters were E(*θ*) = 0, 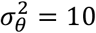, *ω* = 5, *m* = −*ω*/2, *α* = −*m*/5, and *p*_0_ = 10^−3^, and *N*_*e*_ = 10^6^.

### Non-plastic QTL

We first focus on the situation where the phenotypic effect *α* of the mutation at the QTL does not change in response to the environment. The environment is assumed to undergo a sudden shift at time 0 in addition to the stochastic fluctuations, such that the expected mean background phenotype initially deviates from the expected optimum by *d* = E(*m*_0_) − E(*θ*), and that a mutation approaching the mean phenotype from the average optimum is expected to be favored.

#### Monomorphic background

It is informative to first investigate the simplest case where the trait does not have background polygenic variation. The focal mutation at the QTL then segregates in a population that is otherwise monomorphic with respect to adaptation to the fluctuating environment. This context belongs to the weak-mutation limit often assumed in molecular evolution, for instance in Gillespie’s (1983, 1991) SSWM regime, and establishes the most direct connection with results from earlier models of fluctuating selection that do not include an explicit phenotype under selection (Wright 1948; Kimura 1954; Nei 1971; Ohta 1972; Gillespie 1973, 1979, 1991; Nei and Yokoyama 1976; Takahata and Kimura 1979). With monomorphic background, from eq. (13) the expected logit allelic frequency at time *t* starting from a known frequency *p*_0_ is

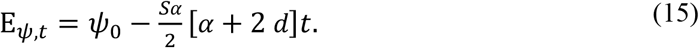

In this context, the expected logit allelic frequency thus increases linearly in time, with a slope given by the expected selection coefficient 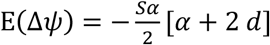. This selection coefficient is not affected by random fluctuations in the optimum, and instead only depends on the constant mismatch *d* between the background mean phenotype *m* and the expected optimum E(*θ*). The mutation at the QTL is expected to spread in the population only if allows approaching the optimum, that is, if *α*^2^ + 2 *αd* < 0.

Even though fluctuations in the optimum do not affect the expected trajectory, they do increase the variance of the stochastic population genetic process. The variance of logit allelic frequency at time *t*, starting from a known frequency *p*_0_, is (from eq. 13),

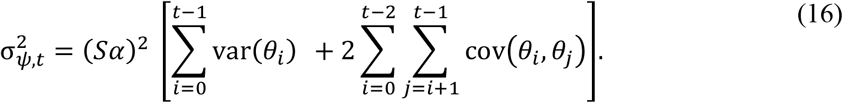

When the optimum undergoes a stationary AR1 process as assumed here, the variance of the population genetic process at the QTL becomes

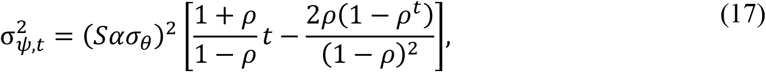

where 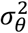 is the stationary variance of random fluctuations in the optimum, and *ρ* is their autocorrelation over one generation. Note that in this scenario, *ρ* is also the per-generation autocorrelation of selection coefficients *s* = Δ*ψ*, while the variance of selection coefficients is Var(Δ*ψ*) = (*Sα*σ_*θ*_)^2^. For large times 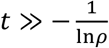, eq. (17) further simplifies as

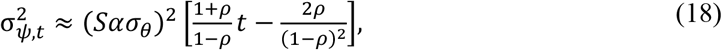

which shows that the variance in logit allelic frequency eventually increases near to linearly with time (**Figure 3A**), and converges more rapidly to this linear change under smaller autocorrelation in the optimum. Stochastic variance in the optimum increases faster under larger autocorrelation in the optimum. **Figure 1** shows that the distribution of *ψ* is well predicted by a Gaussian with mean and variance given by eqs. (15) and (17). Increasing environmental autocorrelation does not change the expected evolutionary trajectory on the logit scale, but increases its variance (**Figure 1**A-B). When transforming to the scale of allelic frequencies, increased environmental autocorrelation causes a broadening of the time span over which selective sweeps occur in the population (**Figure 1**C-D).

##### Polygenic background

With polygenic variation in the background, the mean background phenotype is no longer constant, but instead evolves in response to deterministic and stochastic components of environmental change. Away from the unstable equilibrium in eq. (11), the expected evolutionary trajectory at the QTL is similar to that investigated without fluctuating selection (Lande 1983; Chevin and Hospital 2008). In particular, when the influence of the QTL on evolution of the background trait can be neglected, then combining eqs. (6) and (8) the expected mean background phenotype approaches the expected optimum geometrically, E(*m*) − E(*θ*) = *d*(1 − *SG*)^*t*^ (Lande 1976; Gomulkiewicz and Holt 1995). Combining with eq. (13), the expected logit allelic frequency is

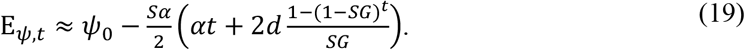

This shows that even when a mutation at the QTL is initially beneficial because it points towards the optimum, its dynamics slows down in time as the mean background approaches the optimum (Lande 1983; Chevin and Hospital 2008). Equation (19) even predicts that an initially beneficial mutation eventually becomes deleterious, and starts declining in frequency when the mean background is sufficiently close to the optimum that the QTL causes an overshoot of the latter (Lande 1983; Chevin and Hospital 2008). This can be seen by noting that in the long run, the term in parenthesis in eq. (19) tends towards *αt* + 2*d*/*SG* and eventually becomes dominated by *αt*, leading to an expected dynamics that declines linearly with slope −*Sα*^2^*t*/2. An initially beneficial mutation starts declining when its selection coefficient crosses 0. Applying the weak-effect approximation for evolution of the mean background (above eq. 19) to eq. (9), this occurs when *α* + 2*d*(1 − *SG*)^*t*^ = 0, that is, at time

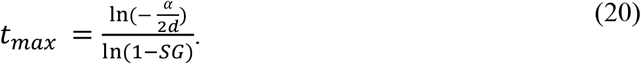

At this point, the expected logit allelic frequency of the mutation at the QTL reaches its maximum, which is (combining eqs. 20 and 19)

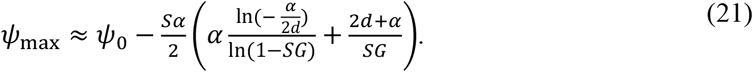

However, this scenario may actually be avoided if the focal mutation reaches *p* > 1/2 (*ψ* > 0) before *t*_*max*_, such that the system gets beyond the unstable equilibrium in eq. (11). The mutation at the QTL then sweeps to fixation, and the mean background evolves away from the optimum to compensate for the QTL effect (Lande 1983; Chevin and Hospital 2008). We will investigate this scenario in more detail below, but let us first turn to the variance of the stochastic process.

For the variance of the process, we rely on the weak-effect approximation in eq. (12), whereby fluctuating selection on the mean background phenotype is little affected by dynamics at the QTL. More broadly speaking, we assume the system is away from the unstable equilibrium in eq. (11). When this holds, we can build upon previous evolutionary quantitative genetics results for the dynamics of the mean background phenotype in a fluctuating environment, to derive the dynamics at the QTL. For an AR1 process as modeled here, the stationary variance of mismatch of the mean background phenotype with the optimum is (Charlesworth 1993)

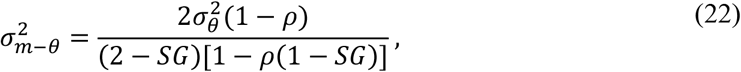

and its temporal autocorrelation function over *τ* generations is

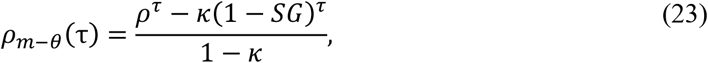

where 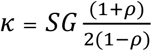 (Cotto and Chevin 2019; see also continuous-time approximation in Chevin and Haller 2014). Combining with eq. (16) leads, after some algebra, to the stochastic variance of logit allelic frequency,

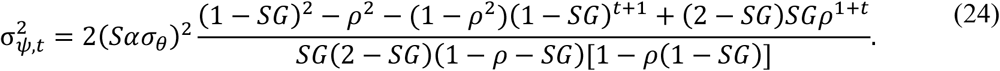

Quite strikingly, contrary to the case of a monomorphic genetic background, 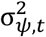 does not increase indefinitely with polygenic background; instead, its dynamics slows down towards an asymptotic maximum,

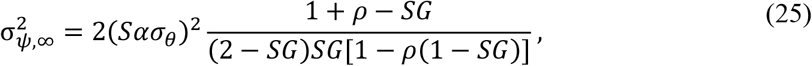

which under weak rate of response to selection in the background *SG* can be approximated by

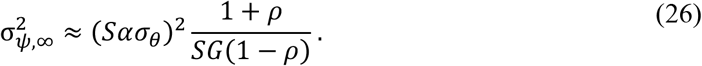

In other words, with a polygenic background, the distribution of logit allelic frequency *ψ* at the QTL tends to a traveling wave, *i.e.* a Gaussian with moving mean but constant variance, as shown in **Figure 2**. This property holds as long as the population is not near the unstable equilibrium in eq. (11), and frequencies at the QTL are sufficiently intermediate that drift is not the main source stochasticity (below).

**Figure 2:**
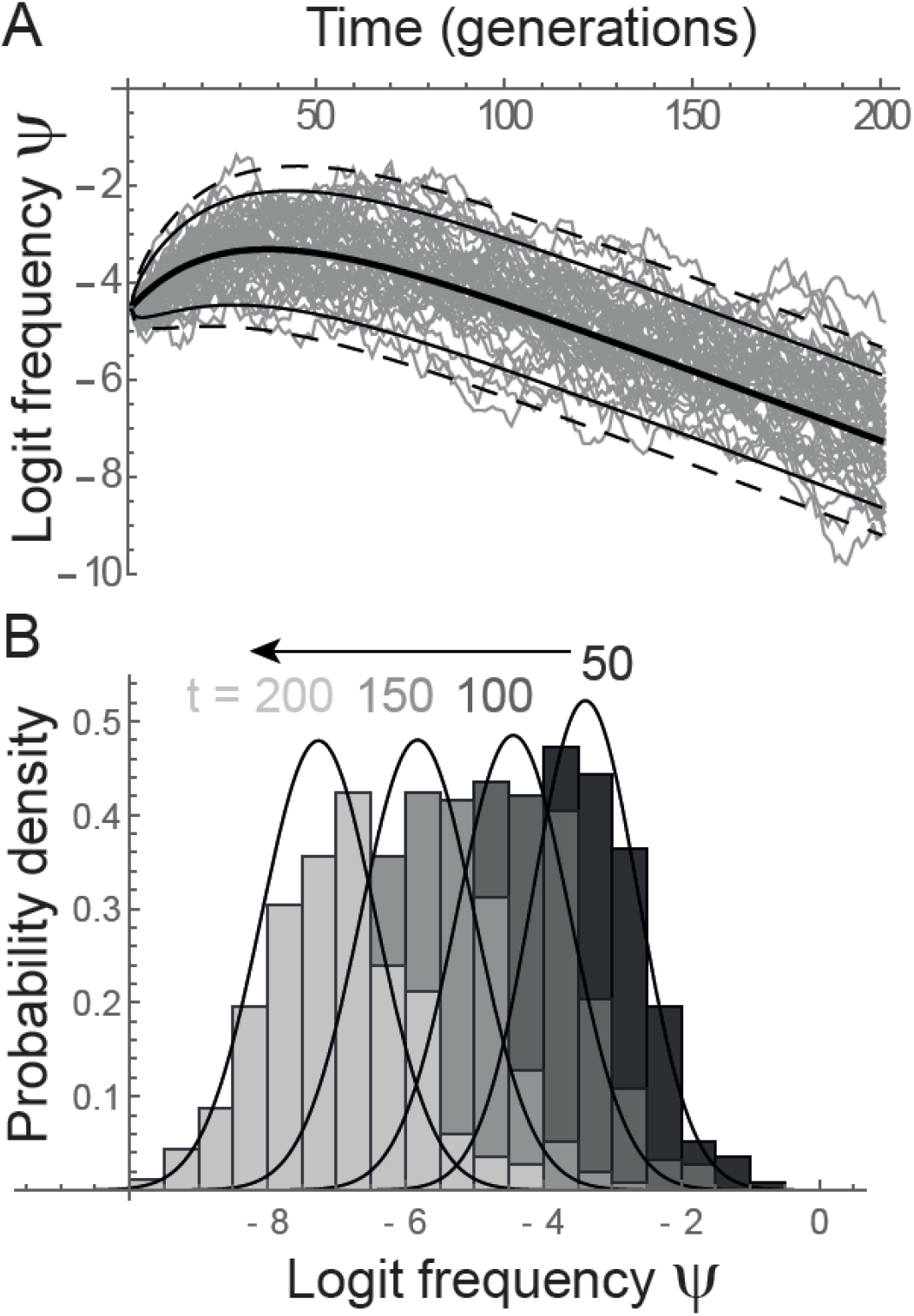
Fluctuating selection at a QTL with a polygenic genetic background. A: The dynamics of logit allelic frequency *ψ* are shown as gray lines for 50 simulations. Also shown are percentiles from the predicted normal distribution, with mean and variance given by eqs. (19) and (24), respectively: 50% (median) in thick, 5% and 95% in thin, and 1% and 99% in dashed lines. B: Histograms show distributions of *ψ* along time for 500 simulations, while the solid black lines are the predicted normal densities based on eqs. (19) and (24). Note how the distribution reaches a stationary variance with a moving mean, that is, a traveling wave with direction given by the arrow in B. Note also that in this example, the sweep at the QTL is interrupted by the mean background evolving towards the optimum, as investigated in detail in **Figure 4**. Parameters were E(*θ*) = 0, 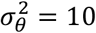, *ρ* = 0.1, *ω* = 5, *m*_0_ = −*ω*/2, *α* = −*m*_0_/2, *G* = 0.5, *p*_0_ = 10^−2^, and *N*_*e*_ = 10^6^.

Inspection of eq. (24) indicates that the rate of approach to the asymptotic variance is determined by the smallest of (1 − *SG*) and *ρ*. In realistic parameter ranges, the rate of response to selection in the background *SG* is small, while *ρ* may be well below 1, so the time scale of approach to equilibrium for 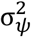 should scale in (*SG*)^−1^. This is confirmed by the simulations, which show that 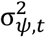 converges faster to its asymptote under larger background genetic variance, while the rate of convergence is little affected by *ρ* (**Figure 3**). The asymptotic variance may be well below that in the absence of polygenic background variation (compare panel A to B-C in **Figure 3**). As predicted by eqs. (25–26), the asymptotic variance 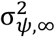 decreases with increasing genetic variance *G* in the background, and increases with increasing environmental autocorrelation *ρ* (**Figure 3**). The influence of autocorrelation is highly non-linear: in our example 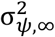 is approximately doubled from *ρ* = 0.1 to *ρ* = 0.5, but multiplied by 4-5 from *ρ* = 0.5 to *ρ* = 0.9 (**Figure 3** B-C).

**Figure 3:**
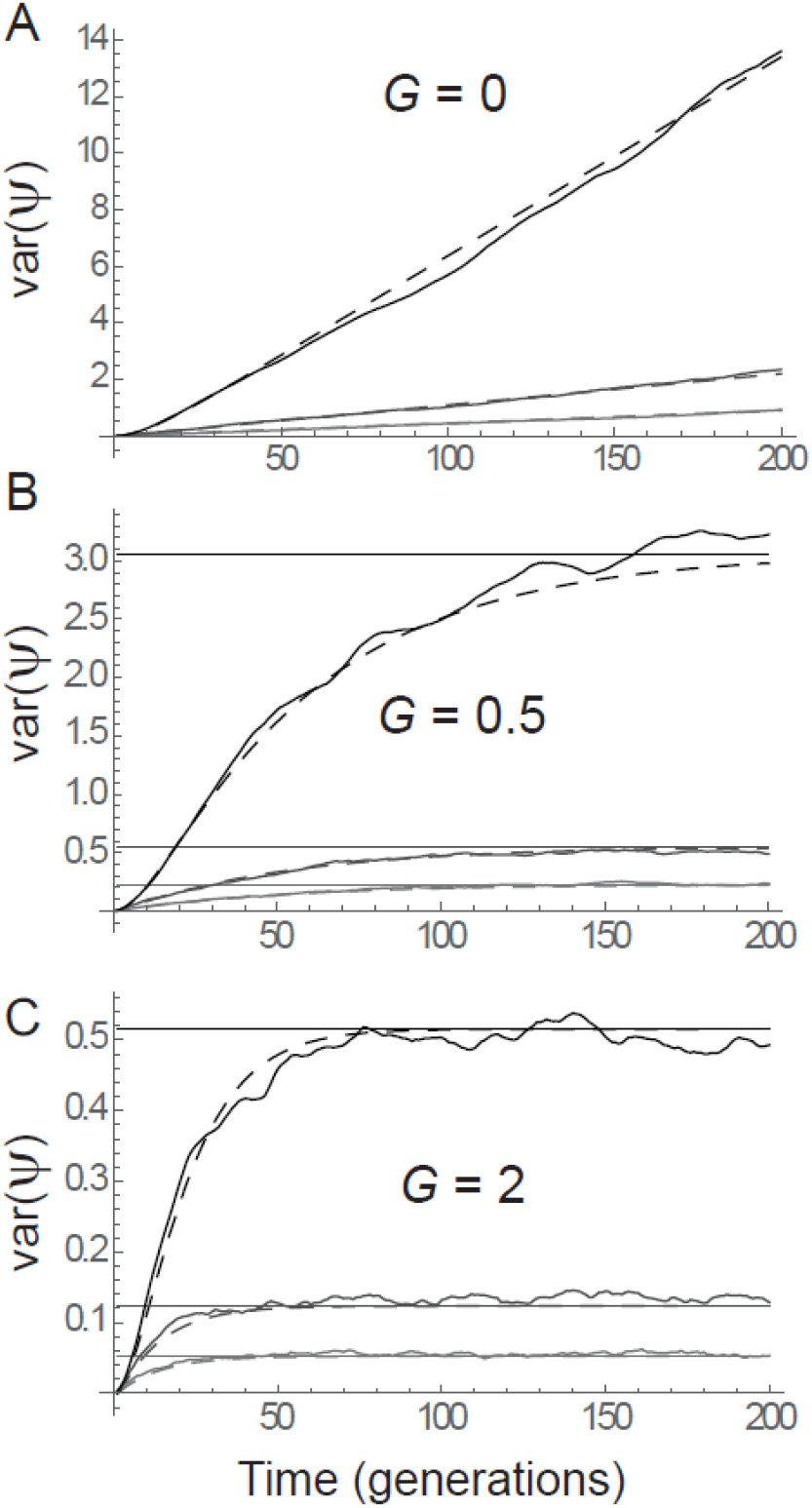
Stochastic variance at the QTL with or without a polygenic background. The variance across replicates of logit allelic frequency *ψ*, starting from a known frequency *p*_0_, is represented along time in simulations without (A) or with (B, C) background polygenic variance for the trait. Also shown in dashed are the predicted dynamics based on equation (17) in A, and eq. (24) in B-C. Note the qualitative difference between the near linear increase in the absence of background genetic variance, versus the saturating dynamics with background genetic variance, for which the maximum asymptotic value from eq. (25) is also plotted as horizontal solid line. The autocorrelation of the optimum is *ρ* = 0.1 (gray), *ρ* = 0.5 (dark gray) or *ρ* = 0.9 (black), additive genetic variance in the background is *G* = 0 (A) *G* = 0.5 (B) or *G* = 2 (C), effective population size is *N*_*e*_ = 10^8^, and other parameters are as in **Figure 1**.

The variance of the stochastic population genetic process has consequences for the bistability of genetic architecture, and the likelihood of a complete sweep. In particular, when the expected trajectory in eq. (19) reaches the vicinity of the unstable equilibrium in eq. (11), the process variance may cause paths to split on each side of this equilibrium and reach alternative fixed equilibria, with either complete sweep or loss of the mutation at the QTL (eq. 10). This is illustrated in **Figure 4**. In this example, the expected trajectory involves a loss of the mutation at the QTL, which occurs for all sample paths shown in **Figure 4**A. However, increasing environmental autocorrelation causes some trajectories to sweep to high frequency (**Figure 4B**). This occurs because environmental autocorrelation increases the stochastic variance of the population genetic process (eqs. 24, 25), and thereby the probability that some trajectories cross the unstable equilibrium, reaching the basin of attraction of the high-frequency equilibrium. Based on this rationale, the proportion of trajectories that reach each alternative stable equilibrium (fixation or loss) may be approximated from the expected proportion of trajectories that are above and below the unstable equilibrium, based on the predicted Gaussian distribution of *ψ* at time *t*_*max*_, when the expected frequency is predicted to be highest based on the simplified model where the QTL does not affect evolution of the mean background (eq. 20). **Figure 4C** shows that this approach correctly predicts how the proportion of sweeps changes with environmental autocorrelation *ρ*. Importantly, since the expected trajectory does not depend on stochastic environmental fluctuations (neither 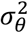 nor *ρ* appear in eq. 19), all effects of environmental autocorrelation (or variance) on the probability of a sweep are mediated by the stochastic variance of the process.

**Figure 4:**
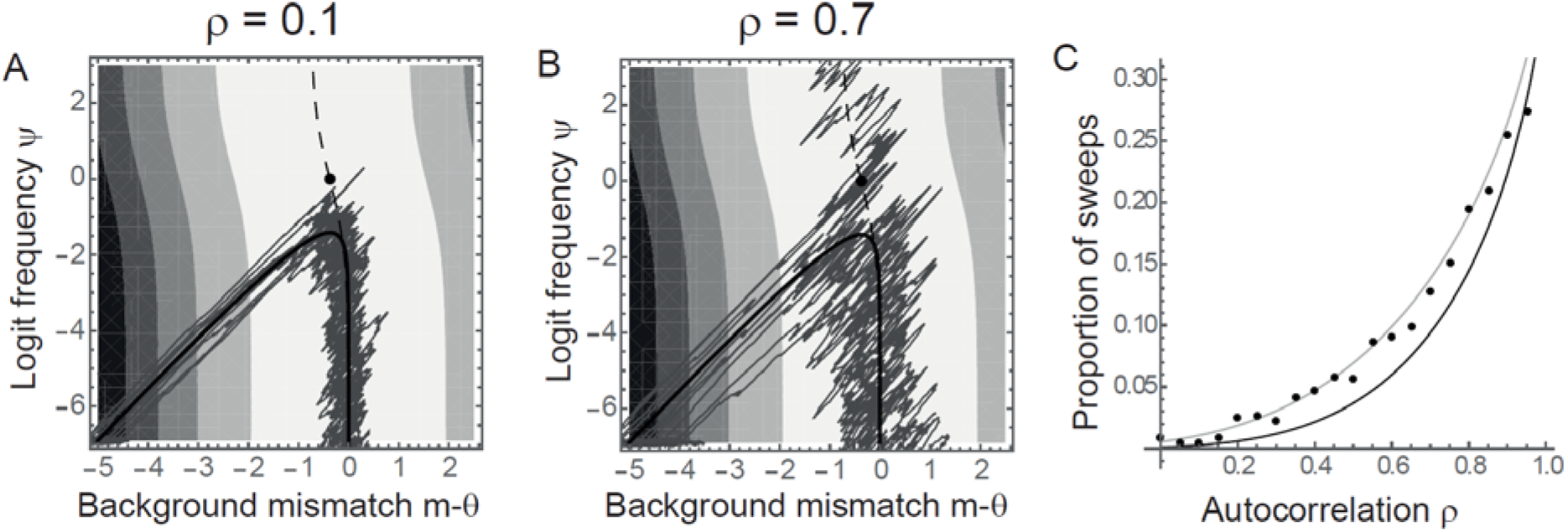
Environmental autocorrelation and probability of a full sweep. The bistability of genetic architecture between major gene and polygenes in this system (eqs. 10–11) is amplified by stochastic fluctuations in the environment. A-B: Joint evolutionary trajectories of logit allelic frequency *ψ* at the major locus and background mean phenotypic deviation from the optimum *m* − *θ* are represented for 10 sampled simulations (dark gray line). The thick black line represents the expected trajectory, neglecting the influence of the QTL on the mean background, obtained by combining eq. (19) with the geometric decline for *m* − *θ*. Shadings represent the fitness landscape in the mean environment, using eq. (7). The dashed line is where the overall mean phenotype is at the optimum, *m* + *pa* = *θ*. All equilibria lie on this line; the unstable equilibrium in eq. (11) is shown as a dot, while the fixed equilibria in eq. 10 cannot be represented on the logit scale. C: The proportion of simulations where the mutation at the QTL eventually reaches frequency higher than 0.95 (dots) is well predicted (lines) using a Gaussian distribution for *ψ*, with equilibrium variance from eq. (25), and mean provided by the expected trajectory at its maximum (eq. 21, black), or the actual maximum frequency in deterministic recursions without environmental fluctuations (gray). For each autocorrelation *ρ* (ranging from 0 to 0.95 by increments of 0.05), 1000 simulations were run, and the proportion of simulations with *p* > 0.95 at generation 2000 was recorded. The parameters for these simulations were *G* = 0.5, E(*θ*) = 0, 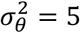, *ω* = 5, *m*_0_ = −*ω*, *α* = −0.15 *m*_0_, and *p*_0_ = 10^−3^, and *N*_*e*_ = 10^6^.

### QTL for phenotypic plasticity

Let us now turn to the case where the QTL influences not only the phenotype, but also how this phenotype responds to the environment. Phenotypic plasticity, the phenotypic response of a given genotype to its environment of development or expression, is a ubiquitous feature across the tree of life (Schlichting and Pigliucci 1998; West-Eberhard 2003). There is also massive evidence for genetic variance in plasticity in the form of genotype-by-environment interactions, one of the oldest and most widespread observations in genetic studies (Falconer 1952; Via and Lande 1985; Scheiner 1993; Gerke *et al.* 2010; Des Marais *et al.* 2013), with molecular mechanisms that are increasingly understood (Angers *et al.* 2010; Beldade *et al.* 2011; Ghalambor *et al.* 2015; Gibert *et al.* 2016). For simplicity I here assume linear reaction norms, where the slope quantifies plasticity. Although this is necessarily a simplification of reality, it is generally a good description over relevant environmental ranges for phenological traits, a major class of phenotypic responses to climate change (e.g., Charmantier *et al.* 2008). This also allows comparing our results to the large body of theoretical literature also based on the assumption of linear reaction norms (Gavrilets and Scheiner 1993b; Scheiner 1998; Lande 2009; Chevin and Lande 2015; Tufto 2015). Such models likely capture the broad evolutionary effects of plasticity for monotonic reaction norms. More complex monotonic reaction norm shapes can be modeled to focus on more specific scenarios such as threshold traits with a bounded range of expression (Chevin and Lande 2013), while non-monotonic reaction norms with an optimum are more appropriate for fitness or performance traits (Lynch and Gabriel 1987; Huey and Kingsolver 1989), which are not the focus here. I also assume for simplicity that the background has constant plasticity, such that all genetic variance in plasticity comes from the major gene. A final assumption in this section will be to focus on stationary environmental fluctuations with no major shift (*d* = 0). Such purely stationary fluctuations are expected to counter-select any mutation at the major gene in the absence of plasticity (eqs. 15 and 19), so it is a good benchmark on which to assess selection on a plasticity QTL.

#### Monomorphic background

In the low mutation limit where the background mean phenotype does not evolve while the mutation is segregating at the QTL, but has still evolved on a longer time scale to match the expected optimum at the onset of selection at the QTL, the expected logit allelic frequency increases linearly in time as in eq. (15), with expected selection coefficient (Appendix)

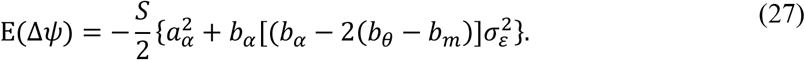

The first term in curly brackets is a component of selection that does not depend on the pattern of environmental fluctuations, and is similar to the expected selection coefficient without plasticity (15) and without a major environmental shift. This component reduces the expected selection coefficient, as it increases the mismatch with the expected optimum phenotype. The second term is a component of selection caused by the effect of the QTL on phenotypic plasticity. This term shows that the plastic effect

*b*_*α*_ of the mutation at the QTL is favored by selection if it allows approaching the optimal response to the environment of development *b*_*θ*_, that is if *b*_*α*_ [(*b*_*α*_ − 2(*b*_*θ*_ − *b*_*m*_)] < 0. The expected selection coefficient is maximum for 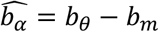, regardless of *a*_*α*_. Importantly, whereas the expected selection coefficient on a non-plastic mutation does not depend on the variance of fluctuations (eq. 15), the component of the expected selection coefficient caused by plasticity is stronger under larger variance 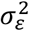 of the environment of development, and thus depends on fluctuations in the optimum (from eq. 4). This reflects the fact that, in a stationary environment, selection on phenotypic plasticity stems from its effect on the variance of phenotypic mismatch with the optimum, rather than on the average mismatch (Lande 2009; Ashander *et al.* 2016). As the variance of the environment of development 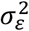 increases, a mutation with a given beneficial effect on phenotypic plasticity becomes increasingly likely to spread even if it causes a systematic mismatch with the optimum in the mean environment, with a deleterious side-effect − 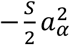. In the absence of background genetic variation, the expected selection at the plasticity QTL does not depend directly on autocorrelation in the environment, but only on the dependence of the optimum on the environment of development, through the parameter *b*_*θ*_. Note however that if phenotypic development/expression and movements of the optimum respond to the same environmental variable (e.g. temperature), but at different times in a generation, then *b*_*θ*_ is directly related to the autocorrelation *ρ* of the optimum (Lande 2009; Michel *et al.* 2014).

The variance of selection coefficients with plasticity but no background genetic variation is

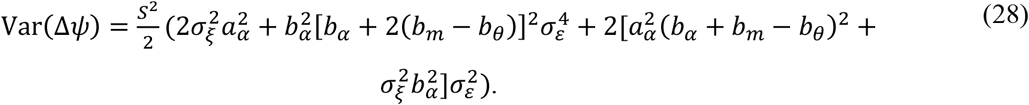

Equation (28) implies that mutations that have the same expected selection coefficient, because they cause the same deviation from the optimal plasticity 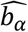, can have different variances in allelic frequency change. This is illustrated in **Figure 5**, which shows that a mutation that leads to hyper-optimal plasticity has more stochastic variance than a mutation that cause equally sub-optimal plasticity, because the former causes overshoots of the optimum while the latter causes undershoots. This difference in stochastic variance between mutations with the same expected selection coefficient, which should impact their relative probabilities of quasi-fixation (Kimura 1954), is stronger for larger deviation from the optimal plasticity (**Figure 5B**).

**Figure 5:**
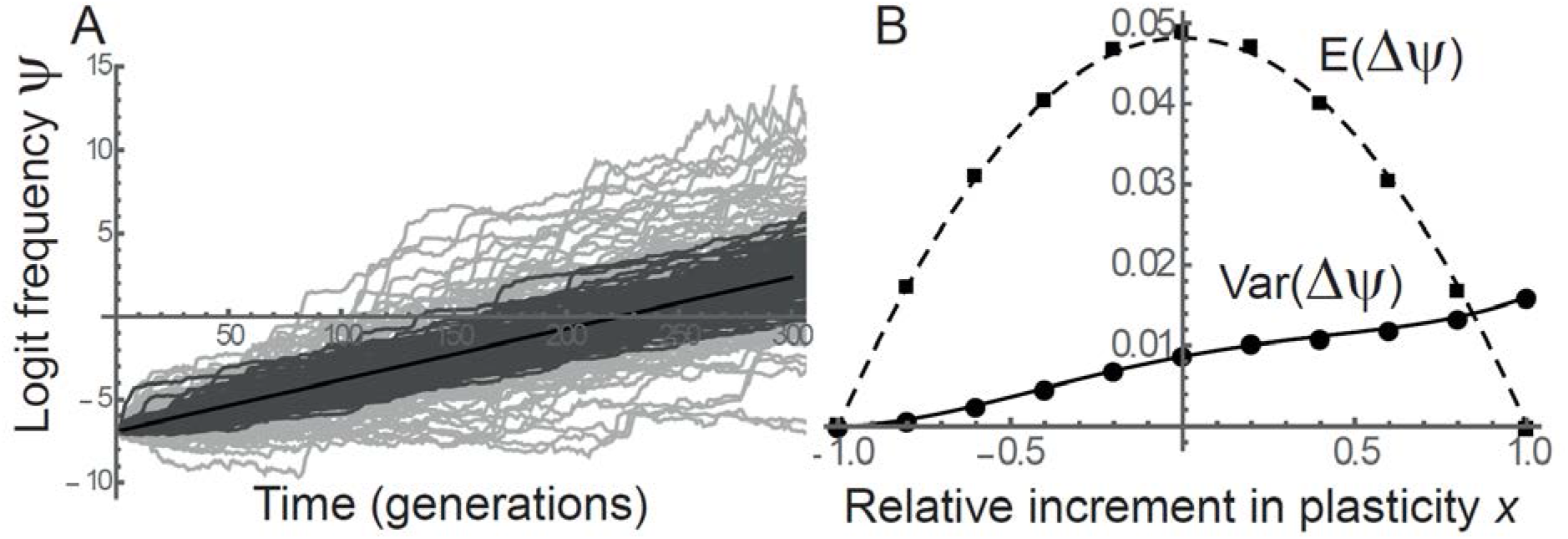
Mean and variance of selection at a QTL for plasticity. A: The dynamics of logit allelic frequency *ψ* at the QTL are represented for simulations where the mutation at the QTL has a small (*b*_*α*_ = 0.4(*b*_*θ*_ − *b*_*m*_), dark gray) or large (*b*_*α*_ = 1.6(*b*_*θ*_ − *b*_*m*_), light gray) effect on phenotypic plasticity, with same expected selection coefficient materialized by the thick black line (based on eq. 27). B: The mean (dashed line: eq. (27); squares: simulations) and variance (continuous line: eq. (28); dots: simulations) of selection coefficients Δ*ψ* are shown as a function of the relative increment *x* in plasticity caused by the mutation at the QTL, such that *b*_*α*_ = (1 + *x*)(*b*_*θ*_ − *b*_*m*_). This shows that mutations with same expected selection coefficient may have different variances in selection, and more so as they deviate more from the optimal plasticity 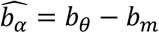 (that is, from *x* = 0). Parameters are 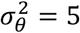, *ρ* = 0.7, 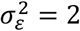, *b*_*θ*_ = 1.4, *b*_*m*_ = 0.2*b*_*θ*_, *a*_*θ*_ = *a*_*m*_ = *a*_*α*_ = 0; other parameters are as in **Figure 1**.

#### Polygenic background

When the mean background phenotype also evolves via polygenic variation, the expected dynamics at the QTL are modified in two main ways. First, background genetic variance contributes to adaptive tracking of the mean phenotype via genetic evolution, thus reducing the benefit of phenotypic plasticity, as in pure quantitative genetic models (Tufto 2015). The level of plasticity that maximizes the expected selection coefficient then becomes (Appendix)

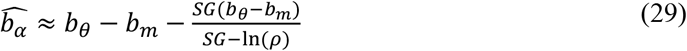

where the last term is the regression slope of the background mean reaction norm intercept on the environment of development, caused by evolution of the mean background in response to the fluctuating environment. **Figure 6A** illustrates how selection via the QTL effect on plasticity is reduced by adaptive tracking of the optimum by evolution of the mean background.

**Figure 6:**
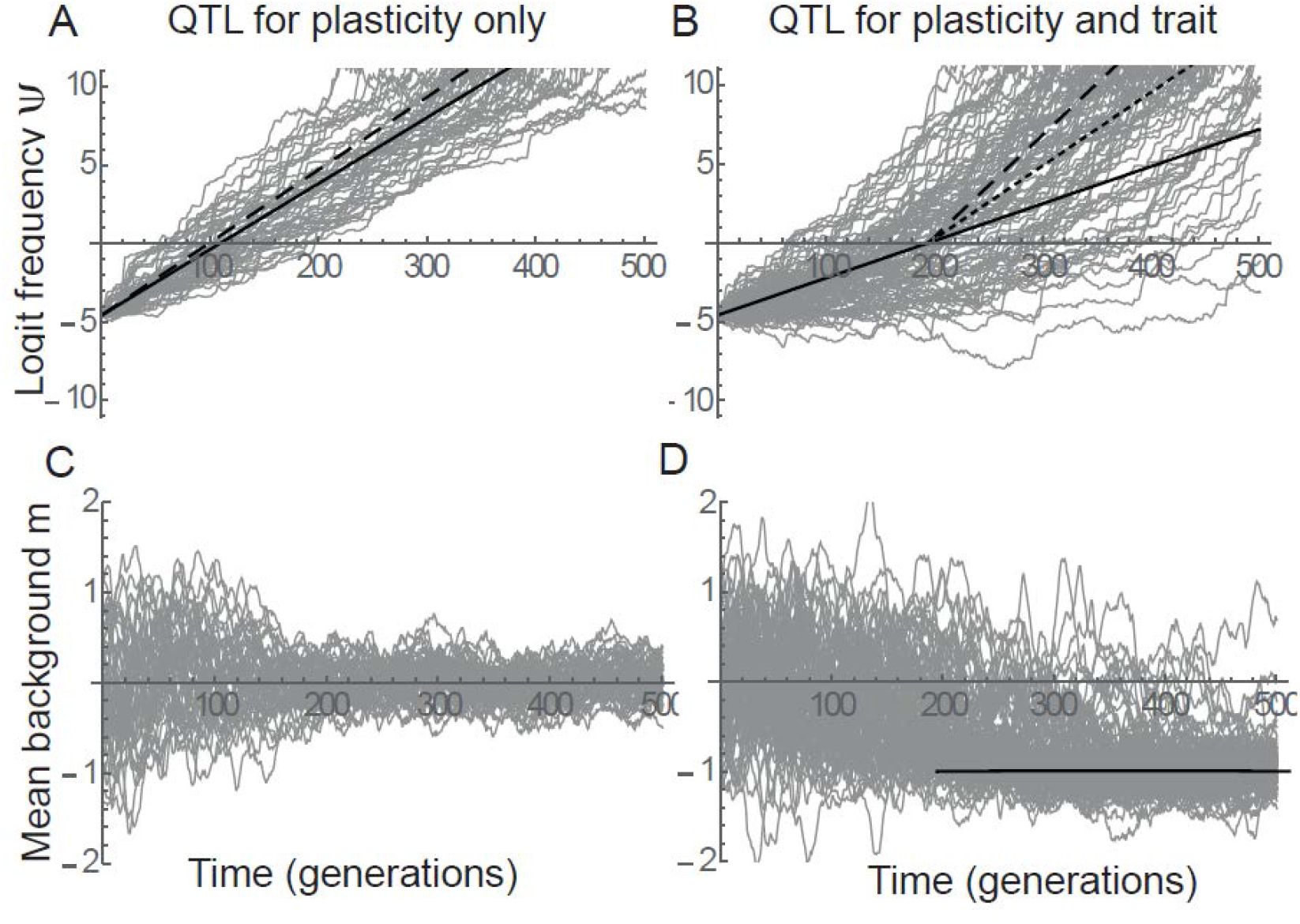
Selection at a QTL for plasticity with background polygenic variation. The dynamics of logit allelic frequency *ψ* at the QTL (A, B) and of the background mean reaction norm elevation *a*_*m*_ (C, D) are represented for the cases where the QTL affects only plasticity (with effect *b*_*α*_ on reaction norm slope, A, C), or also the reaction norm intercept (with effect *a*_*α*_, B, D). In all cases, the gray line show 100 simulations under a randomly changing environment. In panel A, the continuous black line represents the expected dynamics with the selection coefficient in eq. (29), while the dashed line is the prediction that neglects the influence of adaptive tracking of the optimum by the mean background (eq. 27). In panel B, the continuous black line represents the selection coefficient that includes the pleiotropic fitness cost of reaction norm intercept 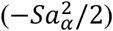, the dashed line represents the selection coefficient that includes a reciprocal benefit 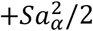, and the dotted line neglect the pleiotropic effect altogether (as in A), after time *t*_*c*_ = ln[(1 − *p*_0_)/*p*_0_]/E(Δ*ψ*). The higher stochastic variance in panel B relative to A is a consequence of the additional effect of the QTL on the reaction norm intercept, consistent with eq (28). In panel D, the black line represents the mean background reaction norm intercept after it has evolved to compensate for the pleiotropic effect of the QTL in the mean environment, such that *a*_*m*_ = *a*_*α*_. Parameters are *G* = 1, *a*_*m*0_ = 0, *a*_*α*_ = 0 (A, C) or *a*_*α*_ = 1 (B, D), *b*_*α*_ = *b*_*θ*_ − *b*_*m*_, *p*_0_ = 10^−2^ and other parameters are as in **Figure 5**.

Second, when the benefit of plasticity allows the mutation at the QTL to spread despite a pleiotropic effect *a*_*α*_ on the intercept of the reaction norm, the expected mean background phenotype can evolve away from the optimum in the average environment to compensate for the associated cost, that is, it evolves to *a*_*m*_ = *a*_*α*_ (**Figure 6D**). Intriguingly, after this has occurred the mutation at the QTL becomes more strongly selected than if it did not have a pleiotropic effect on the reaction norm intercept (**Figure 6B**). This occurs because the QTL effect on reaction norm intercept now allows compensating for maladaptation in the background, which adds a positive component 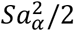 to the benefit *via* the QTL effect on phenotypic plasticity. In other words, what initially caused a displacement from the mean optimum allows approaching the mean optimum after the mean background has been displaced. Furthermore, the spread of the mutation at the plasticity QTL reduces the effective magnitude of fluctuating selection on background mean reaction norm intercept, resulting in smaller evolutionary fluctuations in the background (**Figure 6**C, D).

### Drift versus fluctuating selection

All the analytical results above neglect the influence of random genetic drift, and simulations were run under large *N*_*e*_ to single out the influence of fluctuating selection as a source of stochasticity. However, it is useful to delineate more precisely the conditions under which drift can be neglected relative to environmental stochasticity. The overall variance in allelic frequency change, accounting for both fluctuating selection and random genetic drift in a Wright-Fisher population, can be obtained from the law of total variance, and was previously shown (Ohta 1972) to be

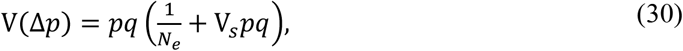

where V_*s*_ = V(Δ*ψ*) is the variance of selection coefficients caused by fluctuating selection. From this it entails that fluctuation selection dominates drift as a source of stochasticity when 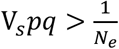, that is for

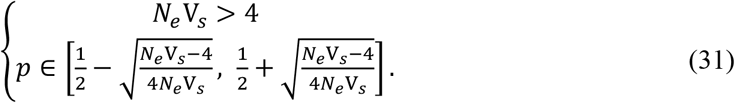

This can be translated into a condition for the logit allelic frequency *ψ*,

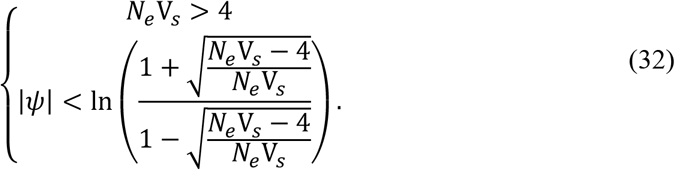

Very similar results are obtained (not shown) if the criterion is based on the stochastic variance of *ψ*, for which the fluctuating selection component is independent of *ψ* (as derived in the main text), but the drift component is not. Equation (32) shows that an absolute condition for fluctuating selection to be the dominant source of stochasticity is *N*_*e*_V_*s*_ > 4. When this holds, fluctuating selection dominates over a range of intermediate allelic frequencies, while drift dominates at extreme frequencies outside of this range. The bounds of this range are entirely determined by the compound parameter *N*_*e*_ V_*s*_, as shown by eqs. (31-32) and **Figure 7A**. **Figure 7** further illustrates that for small *N*_*e*_ V_*s*_, small initial frequencies and/or large final frequencies result in inflated variance relative to the expectation under pure fluctuating selection (panels B-C), as well as fixation events by drift (panel B). As *N*_*e*_ V_*s*_ increases from panel B to D, the predictions under pure fluctuating selection become increasingly accurate, all the more so as the initial allelic frequency is within the range defined by eq. (32).

**Figure 7:**
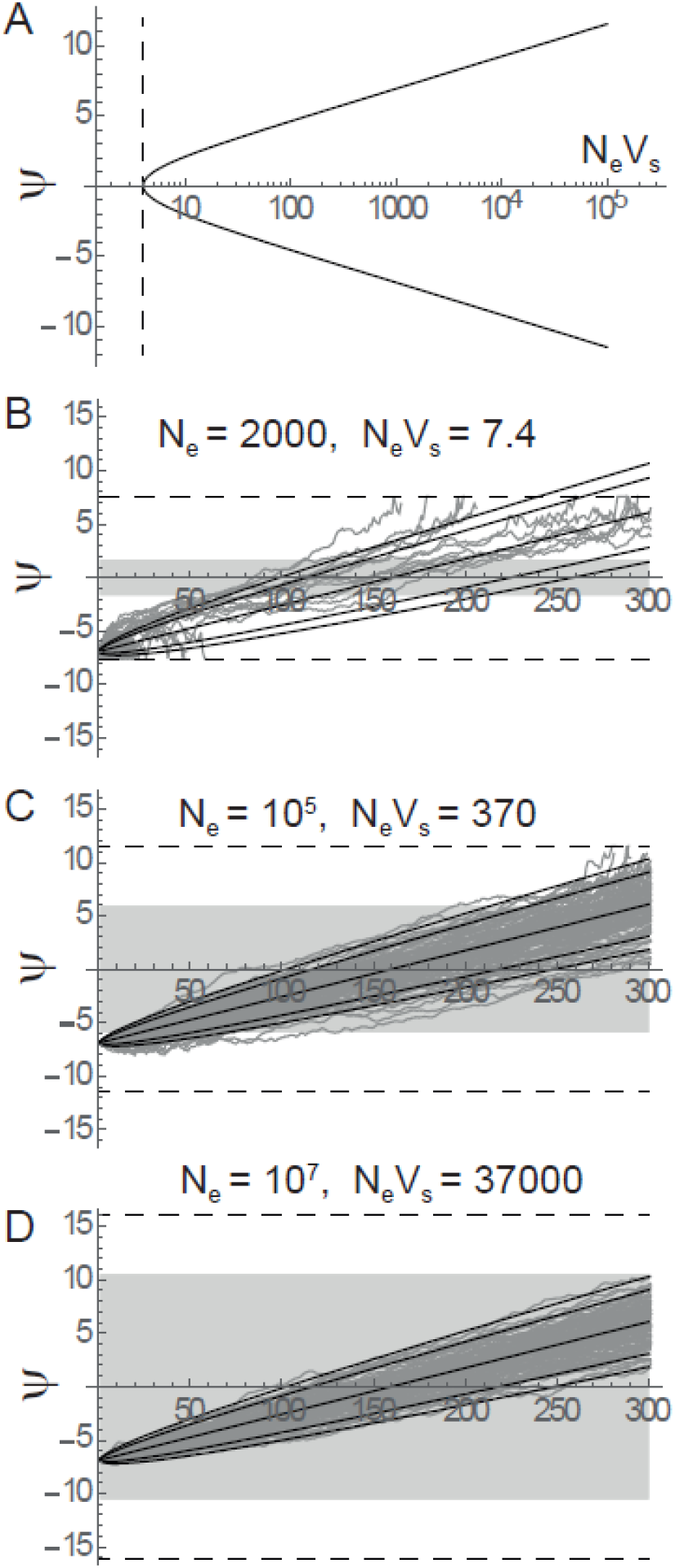
Drift versus fluctuating selection. A: The threshold logit frequency beyond which drift dominates fluctuating selection as a source of stochasticity (from eq. 32, full line) is plotted against the compound parameter *N*_*e*_ V_*s*_. The dashed line represents the hard threshold at *N*_*e*_V_*s*_ = 4. B-D: The dynamics of logit allelic frequency *ψ* is plotted over time for 50 simulations with selection and drift, and without plasticity or background genetic variation, similarly to **Figure 1**. The continuous lines show the predicted quantile of the distribution, as in **Figure 1**. The shaded region indicates the range of *ψ* over which fluctuating selection is expected to dominate, using eq. (32) with V_*s*_ = (*Sα*σ_*θ*_)^2^. The dashed lines show the fixation threshold at ± ln *N*_*e*_. The effective population sizes are indicated in each panel, and other parameters are as in **Figure 1**.

## Discussion

Analysis of a simple model combining population and quantitative genetics has revealed a number of interesting properties about fluctuating selection at a gene affecting a quantitative trait (or QTL), when this trait undergoes randomly fluctuating selection caused by a moving optimum phenotype. The first important observation is that, when assessed on the logit scale - a natural scale for allelic frequencies (Kimura 1954; Gillespie 1991; Chevin 2011; Gallet *et al.* 2012) -, the dynamics at the QTL has a simple connection to movements of the optimum, since the selection coefficient depends linearly on the mismatch between the mean background phenotype and the optimum (eq. 9; see also Martin and Lenormand 2006). For a QTL that has the same phenotypic effect in all environments (no phenotypic plasticity), the expected trajectory only depends on the expected phenotypic mismatch with the optimum, not on the pattern of fluctuations in this optimum. However the variance of trajectories, an important determinant of probabilities of quasi-fixation (Kimura 1954), is strongly affected not only by the magnitude of fluctuations in the optimum, but also by their autocorrelation (eq. 17, **Figure 1**). When the focal QTL is the only polymorphic gene undergoing fluctuating selection, this stochastic variance increases linearly over time (**Figure 3A**), at a rate that is faster under larger positive autocorrelation in the optimum. In contrast, when polygenic variation elsewhere in the genome allows for evolution of the mean background phenotype, stochastic variance at the QTL is bounded by a maximum asymptotic value, which is lower under higher genetic variance in the background (eqs. 24–25 and **Figure 3**B-C). This stochastic variance caused by fluctuating selection interacts with the inherent bi-stability of genetic architecture in this system (Lande 1983; Chevin and Hospital 2008), and may increase the probability that the mutation at the QTL reaches fixation at the expense of the background mean phenotype (as illustrated in **Figure 4**), or the reverse.

When the mutation at the QTL also affects phenotypic plasticity via the slope of a linear reaction norm, then even its *expected* trajectory depends on the pattern of fluctuations, with stronger selection under large fluctuations (eq. 27), contrary to the case of a non-plastic QTL. Interestingly, mutations with the same expected selection coefficient - because they cause the same deviation from the optimal plasticity – may have very different variances in allelic trajectories, depending on whether they tend to cause overshoots or undershoots of the fluctuating optimum (**Figure 5**). Finally, a mutation that is sufficiently strongly selected via its effect on phenotypic plasticity can spread despite causing a systematic mismatch with the optimum in the average environment. When the mean background phenotype can evolve by polygenic variation, it can compensate for this pleiotropic effect on reaction norm intercept. Quite strikingly, this increases selection at the plasticity QTL, causing the mutation to spread faster than if it only affected plasticity (**Figure 6B**).

Consistent with previous uses of this model with a major gene and polygenes (Lande 1983; Agrawal *et al.* 2001; Chevin and Hospital 2008; Gomulkiewicz *et al.* 2010), I did not model explicitly the maintenance of genetic variance in the background, instead assuming that it had reached an equilibrium between mutation and stabilizing/fluctuating selection. This has provided simple and robust analytical insights about the interplay of selection at a major gene with background polygenic variation. Although environmental fluctuations should affect the expected additive genetic variance *G* to some extent (Burger and Gimelfarb 2004; Svardal *et al.* 2011), this does not necessarily affect our results because they are conditioned on *G*, rather than on mutational variance for instance, which is less directly amenable to empirical measurement. More critical is the fact that the background additive genetic variance should itself fluctuate in time as alleles in the background change in frequency, especially in a finite population (Bürger and Lande 1994; Höllinger *et al.* 2019). This should increase temporal variation in the evolutionary process, so that results about stochastic variance here may be considered as lower bounds, if the long-term mean *G* is used in formula. Modeling explicitly the dynamics of background quantitative genetic variance in a random environment would require using individual-based simulations, as done for instance by Bürger and Gimelfarb (2002). Previous work based on a similar environmental context as modeled here proved that most results are little affected in regimes where substantial genetic variation can be maintained for a quantitative trait (Chevin and Haller 2014; Chevin *et al.* 2017), as assumed here.

Although the simulations included random genetic drift, all the analytical results were derived by neglecting the influence of drift. These analytical results are therefore valid over a range of allelic frequencies that is entirely determined by the product of the effective size by the variance of selection coefficients, as shown in eqs. (31–32) and **Figure 7**. In most simulations, I have assumed that the mutation at the QTL is initially at low frequency, but still common enough to be within the range defined by eqs. (31–32), where frequency change is entirely driven by selection. It would be worthwhile investigating in future work the probability of establishment of a mutation that starts in one copy and affects a trait exposed to randomly fluctuating selection, but this requires developments that are beyond the scope of the present study. For our purpose, we can consider that the initial frequency *p*_0_ stems either from the trajectory of a newly arisen mutation conditional on non-extinction, which is expected to rapidly rise away from 0 (Barton 1998; Martin and Lambert 2015), or from a distribution at mutation-selection drift equilibrium (Wright 1937; Barton 1989; Höllinger *et al.* 2019).

Our analytical results about the distribution of logit allelic frequency lend themselves well to comparisons with empirical measurements. Indeed the logit of allelic frequencies is readily obtained from number of copies of each type, since 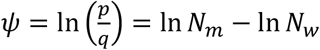, where *N*_*m*_ and *N*_*w*_ are the copy numbers of the mutant (derived) and wild-type (ancestral) allele, respectively. In fact, when frequencies are estimated on a subsample from the population, the strength of selection on genotypes is generally estimated using logistic regression (Gallet *et al.* 2012), a generalized linear (mixed) model that uses the logit as link function. Our theoretical predictions therefore apply directly to the linear predictor of such a GLMM, without requiring any transformation. For instance if we consider an experiment where multiple lines undergo independent times series of a stochastic environment (*i.e.*, different paths of the same process), the stochastic variance among replicates can be estimated as a random effect in a logistic GLMM. If multiple loci are available, this random effect should strongly covary among loci within an environmental time series, because they share the same history of environments, in contrast to frequency changes caused by drift, which should only be similar between tightly linked loci.

The results here are based on a model of fluctuating optimum for a quantitative trait, similar to previous theory by Connallon and Clark (2015), but extend this theory by allowing for environmental autocorrelation, and by deriving the stochastic variance of the population genetic process. Importantly, most of the present results should also be relevant to cases where an explicit phenotype under selection is not identified or measured, but the relationship between fitness and the environment has the form of a function with an optimum, which can be approximated as Gaussian (Lynch and Gabriel 1987; Gabriel and Lynch 1992; Gilchrist 1995). For many organisms, especially microbes, measuring individual phenotypes can be challenging, and it may prove difficult to identify most traits involved in adaptation to a particular type of environmental change (ie temperature, salinity…). A common solution is to directly measure fitness or its life-history components (survival, fecundity) across environments, to produce an environmental tolerance curve (Deutsch *et al.* 2008; Thomas *et al.* 2012; Foray *et al.* 2014). An influence of the history of previous environments on these tolerance curves can also be included, via plasticity-mediated acclimation effects (Calosi *et al.* 2008; Gunderson and Stillman 2015; Nougué *et al.* 2016). It has been highlighted previously that tolerance curves can be thought of as emerging from a moving optimum phenotype on unmeasured, possibly plastic, underlying traits (Chevin *et al.* 2010; Lande 2014), so that a simple re-parameterization can translate all the results above in terms of evolution of tolerance breadth and environmental optimum. Such a connection has recently been invoked to analyze population dynamics in a stochastic environment (Chevin *et al.* 2017; Rescan *et al.* 2019), suggesting that results from the current study are not restricted to cases where relevant quantitative traits under fluctuating selection can be measured, but may instead apply to a broad range of organisms exposed to randomly changing environments.

## Acknowledgements

This work was supported by the European Reseach Council (Grant 678140-FluctEvol). I thank J. Hermisson and three anonymous reviewers for useful criticisms and suggestions.

## Appendix Details of plasticity model

In the model with plasticity, the environment is assumed to undergo stationary fluctuations, before and after the appearance and spread of the mutation at the QTL. Before the mutation at the QTL reaches appreciable frequency, the recursion for the mean background phenotype is (combining eqs. (2), (4), (6) and (8))

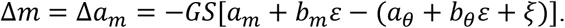

Integrating over the distribution of environments of development *ɛ* and residual component of variance in the optimum *ξ*, the expected mean reaction norm intercept at equilibrium in a stationary environment, before the mutation at the QTL establishes and starts spreading, is

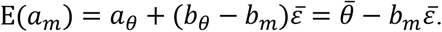

This shows that the mean reaction norm intercept evolves so as to compensate for the effect of plasticity, such that the overall mean background phenotype 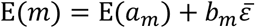 is at the expected optimum 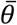. However, the intercept of a reaction norm has no meaning per se, as it depends on the arbitrary choice of a reference environment where *ɛ* = 0. We thus choose to set as reference the stationary mean of the environment of development, *de facto* setting 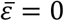. This is just a way of parameterizing the model such that the intercept for the optimum is simply the stationary mean optimum, 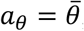, which is also equal to the expected reaction norm intercept E(*a*_*m*_) in the absence of any influence from the QTL.

The recursion for the change in logit allelic frequency over a generation can be obtained by combining equations (9) and (2–4), leading to

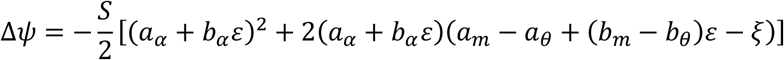

which can be expanded to yield

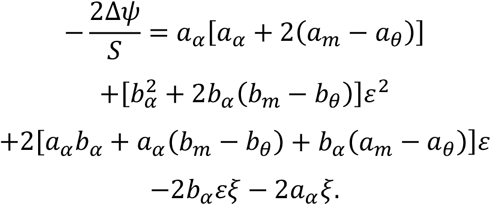

Integrating over the distribution of environments of development *ɛ* and residual component of variance in the optimum *ξ* yields

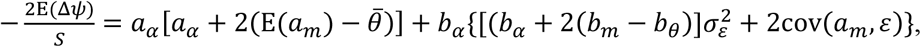

where the covariance cov(*a*_*m*_, *ɛ*) is caused by adaptive tracking of the moving optimum phenotype by evolution of the mean background phenotype. In the absence of polygenic variation during the sweep, we have 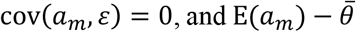 in the long run in a stationary environment, leading to eq. (27) in the main text. With polygenic variance in the background, we have, from Michel et al. (2014),

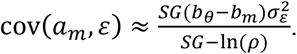

The stochastic variance of logit frequency change is

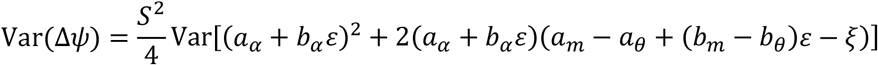

In the absence of background genetic variance, we have

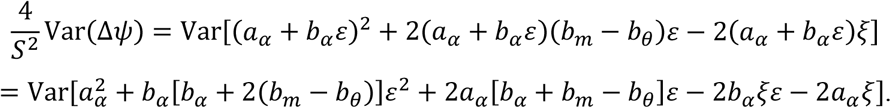

Integrating over the distribution of environments of development *ɛ* and residual component of variance in the optimum *ξ*, this yields

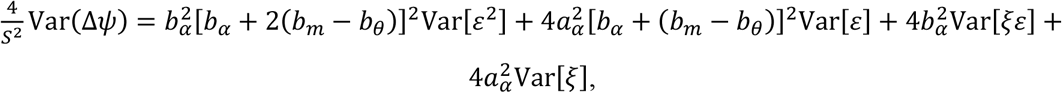

where we have used the fact that, when *ɛ* and *ξ* are independent and with mean 0,

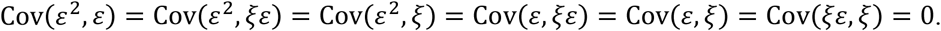

We can also use

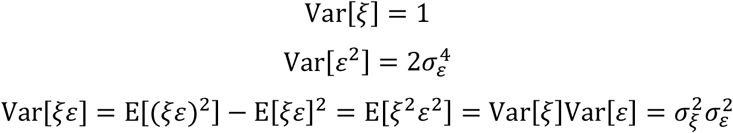

To get

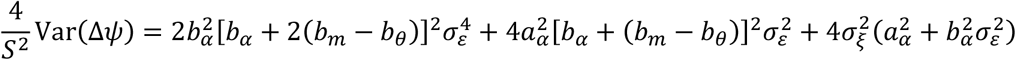

such that

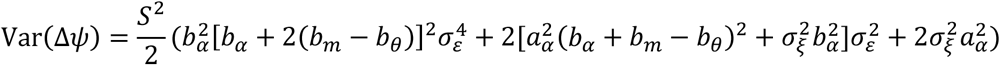

In the simpler case where the mutation only affects plasticity, but not the reaction norm intercept, this simplifies as

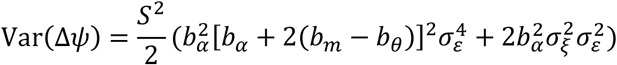

